# Rest/Stress Myocardial Perfusion Imaging by Positron Emission Tomography with ^18^F-Flurpiridaz: A Feasibility Study in Mice

**DOI:** 10.1101/2021.09.05.459034

**Authors:** Susan Bengs, Geoffrey I. Warnock, Angela Portmann, Nidaa Mikail, Alexia Rossi, Hazem Ahmed, Dominik Etter, Livio Gisler, Caitlin Jie, Alexander Meisel, Claudia Keller, Roger Schibli, Linjing Mu, Ronny R. Buechel, Philipp A. Kaufmann, Simon M. Ametamey, Catherine Gebhard, Ahmed Haider

## Abstract

**Background:** Myocardial perfusion imaging by positron emission tomography (PET-MPI) is the current gold standard for quantification of myocardial blood flow. ^18^F-flurpiridaz was recently introduced as a valid alternative to currently used PET-MPI probes. Nonetheless, optimum scan duration and time interval for image analysis are currently unknown. Further, it is unclear whether rest/stress PET-MPI with ^18^F-flurpiridaz is feasible in mice.

**Methods:** Rest/stress PET-MPI was performed with ^18^F-flurpiridaz (0.6-3.0 MBq) in 29 mice aged 7-8 months. Regadenoson (0.1 μg/g) was used for induction of vasodilator stress. Kinetic modeling was performed using a metabolite-corrected arterial input function. Image-derived myocardial ^18^F-flurpiridaz uptake was assessed for different time intervals by placing a volume of interest in the left ventricular myocardium.

**Results:** Tracer kinetics were best described by a two-tissue compartment model. *K*_*1*_ ranged from 6.7-20.0 mL/cm^3^/min, while myocardial volumes of distribution (*V*_*T*_) were between 34.6 and 83.6 mL/cm^3^. Of note, myocardial ^18^F-flurpiridaz uptake (%ID/g) was significantly correlated with *K*_*1*_ at rest and following pharmacological stress testing for all time intervals assessed. However, while Spearman’s coefficients (r_s_) ranged between 0.478 and 0.672, R^2^ values were generally low. In contrast, an excellent correlation of myocardial ^18^F-flurpiridaz uptake with *V*_*T*_ was obtained, particularly when employing the averaged myocardial uptake from 20-40 min post tracer injection (R^2^ ≥0.98). Notably, *K*_*1*_ and *V*_*T*_ were similarly sensitive to pharmacological stress induction. Further, mean stress-to-rest ratios of *K*_*1*_, *V*_*T*_, and %ID/g ^18^F-flurpiridaz were virtually identical, suggesting that %ID/g ^18^F-flurpiridaz can be used to estimate CFR in mice.

**Conclusion:** Our findings suggest that a simplified assessment of relative myocardial perfusion and coronary flow reserve (CFR), based on image-derived tracer uptake, is feasible with ^18^F-flurpiridaz in mice, enabling high-throughput mechanistic CFR studies in rodents.

## Introduction

In the past two decades, the diagnosis of coronary artery disease (CAD) has undergone a remarkable evolution from conventional anatomical assessment of coronary arteries to an integral diagnostic approach combining anatomical and functional cardiovascular imaging modalities. Although coronary computed tomography angiography (CCTA) remains the first-line diagnostic tool for patients with suspected CAD [1], a significant proportion of patients presenting with angina show no evidence of epicardial stenosis [2] and may suffer from a condition called INOCA (ischemia with no obstructive coronary arteries) caused by vasospastic disorders or microvascular dysfunction [3]. The latter is associated with a reduced coronary flow reserve (CFR) and can be quantified by rest/stress myocardial perfusion imaging (MPI) with positron emission tomography (PET). As such, PET-MPI has shown a high diagnostic accuracy and independent prognostic value in INOCA patients [4].

Given the availability of rubidium-82 generators, which obviates the need for a cyclotron, rubidium-82 is the most frequently used probe for PET-MPI. Nonetheless, the relatively high positron range results in an inferior spatial resolution, as compared to other PET isotopes [5]. In contrast, ^15^O-water and ^13^N-ammonia exhibit superior spatial resolutions to rubidium-82 and are validated perfusion tracers, however, their short physical half-lives do not allow satellite distribution, thus, rendering an on-site cyclotron compulsory [6]. Efforts to develop a radiofluorinated myocardial perfusion tracer with increased physical half-life and spatial resolution have ultimately led to the discovery of ^18^F-flurpiridaz [7–9]. Unlike other probes for PET-MPI, ^18^F-flurpiridaz binds to the mitochondrial complex I (MC-I), which is highly abundant in cardiomyocytes [6, 9]. The latter allows selective visualization of the myocardium, providing remarkably high signal-to-background ratios and thus high image quality [10–16]. In experimental models, ^18^F-flurpiridaz exhibited a relatively high myocardial extraction fraction of 0.94, which was consistent at different flow rates, and demonstrated favorable performance characteristics as a myocardial perfusion tracer [8, 9, 17–22].

Despite the abovementioned advantages of ^18^F-flurpiridaz, the physical half-life of 109.8 minutes constitutes a major challenge for efficient rest/stress testing protocols. Indeed, conventional 1-day protocols, as they are routinely performed with rubidium-82, ^13^N-ammonia or ^15^O-water, are hampered by the residual myocardial radioactivity from the initial ^18^F-flurpiridaz injection, which affects the time-activity curves (TACs) obtained after the second ^18^F-flurpiridaz injection. To overcome this limitation, a 2-day protocol has recently been suggested, which allows complete washout of myocardial radioactivity between rest and stress scans [18]. Nonetheless, 2-day protocols are demanding for the patient and less cost-effective than 1-day protocols. Further, several unanswered questions remain regarding the ideal scan duration with ^18^F-flurpiridaz, as well as the most accurate time frame post injection for the assessment of myocardial blood flow (MBF) and CFR. Accordingly, we sought to (1) identify the optimal time frame for PET imaging following ^18^F-flurpiridaz injection and (2) to assess whether a simplified model would allow sequential rest and stress PET-MPI, thereby providing accurate information on MBF and CFR in mice.

## Methods

### Animals

Animal care and experimental procedures were performed in accordance with the Swiss Animal Welfare legislation and approved by the Veterinary Office of the Canton Zurich, Switzerland. Female and male FVB/N mice (N=29, 14 females) were obtained from Janvier Labs (Le Genest-Saint-Isle, France), kept with free access to food and water and were scanned at the age of 7-8 months.

### Study protocol and image acquisition

^18^F-Flurpiridaz radiolabeling and precursor synthesis were reported elsewhere [23–25]. PET/CT was performed in animals that were anaesthetized using 1.3-2.0% isoflurane in oxygen-enriched air (1:1). Depth of anesthesia was monitored via respiratory rate measurement (SA Instruments, Inc., Stony Brook, USA). Body temperature was monitored using a rectal probe and was kept at 37°C with a temperature-adjusted air stream. A dose of 0.6-3.0 MBq ^18^F-flurpiridaz was administered via tail-vein injection 60 sec after start of the PET scan. Tracer distribution was recorded in dynamic PET acquisition mode over a time period of 41 min, before a bolus of regadenoson (0.1 μg/g) and a second dose of ^18^F-flurpiridaz (2.2-8.1 MBq) were injected via a pre-installed intravenous catheter. PET imaging was performed with a calibrated Super Argus PET/CT scanner (Sedecal, Spain), followed by a CT scan for anatomical information. PET data reconstruction was carried out using the manufacturer’s 2D iterative (OSEM, ordered subset expectation maximisation, 2 iterations, 16 subsets) algorithm, and corrections for dead time, decay, scatter and attenuation, at a voxel size of 0.3875 × 0.3875 × 0.775 mm^3^. All radioactivities were decay-corrected to the time of tracer injection.

### Image analysis and generation of rest/stress time-activity curves

Reconstructed PET data were processed with PMOD v.3.8 (PMOD Technologies Ltd., Switzerland) via manual delineation of the myocardium to generate volumes of interest (VOIs) and respective time-activity curves. Decay corrected time-activity curves were calculated for the myocardium VOI in either as kBq/cc or % injected dose per gram tissue (%ID/g). An exponential model was fitted for each rest scan to estimate the radioactivity that remained in the myocardium at 41 to 82 min post injection from the initial ^18^F-flurpiridaz injection (**Figure 1A,** extrapolation curve). To obtain the corrected time-activity curves following stress injection (**Figure 1A**, ^18^F-flurpiridaz and regadenoson), extrapolation curves were subtracted from the respective image-derived time-activity curves at 41 to 82 min post injection. Correction for the injected dose yielded the final rest and stress time-activity curves (**Figure 1B**).

**Figure 1:**
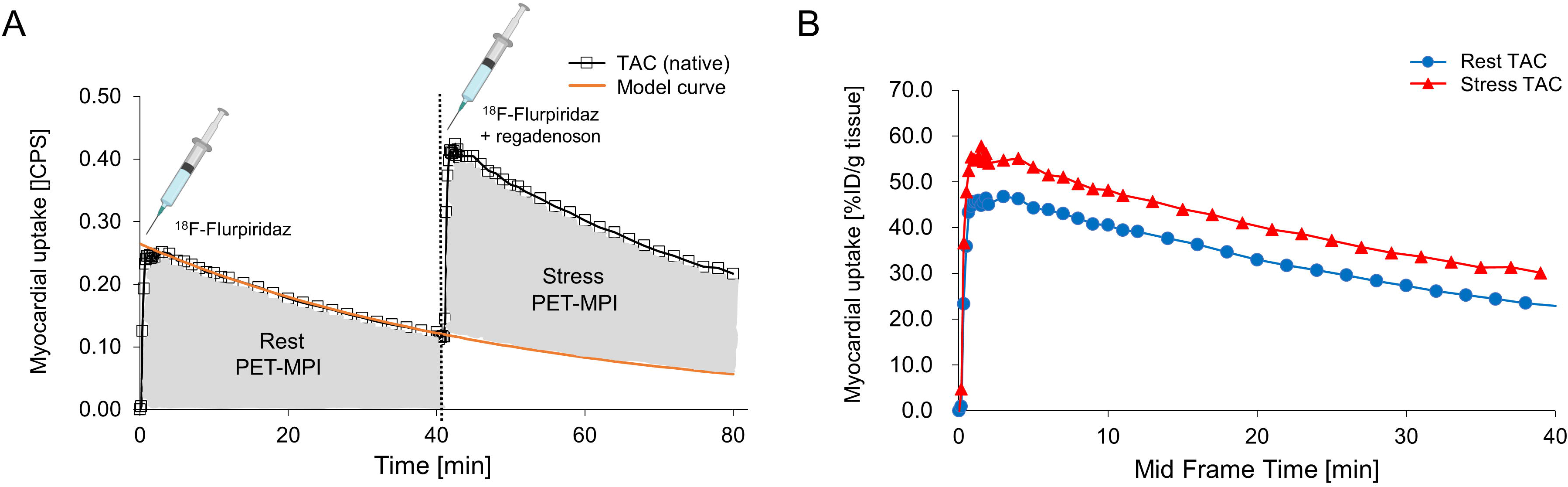
Time-activity curves (TACs) of the mouse myocardium upon tail vein injections of ^18^F-flurpiridaz. **A**. Native TACs after rest (^18^F-flurpiridaz) and stress (^18^F-flurpiridaz + regadenoson) injection, presented as counts per second (CPS). The model curve was fitted with a single exponential model to subtract ^18^F-flurpiridaz injection 1 from injection 2. **B**. Final rest and stress TACs after correction for injected ^18^F-flurpiridaz dose, presented as percent injected dose per gram tissue (%ID/g).

### Input function

An arteriovenous shunt and portable coincidence detector (twilite, swisstrace, Menzingen, Switzerland) were used to record the tracer concentration in blood simultaneously with the PET data acquisition in five FVB/N mice (30.1–32.6 g). The coincidence detector was cross-calibrated to the PET for kBq/cc output. The delay between tissue and blood time-activity curves was determined as part of the model fitting procedure and the final input function was corrected for plasma metabolites and plasma-to-blood ratio (supplemental information). The resulting five input functions were averaged, and the average was used as a surrogate input function for all animals scanned without arteriovenous shunt system (supplemental information).

### Kinetic modeling

Kinetic modeling was performed with the PKIN tool of PMOD v.3.8 (PMOD Technologies Ltd., Switzerland). The kinetics of ^18^F-flurpiridaz was evaluated with a one- and two-tissue compartment model using the following equations:

One-tissue compartment model:

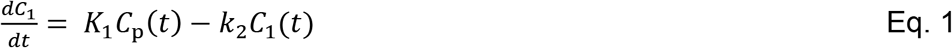

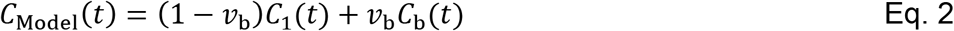

Two-tissue compartment model:

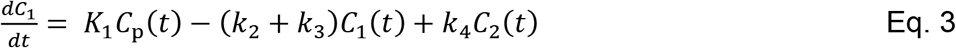

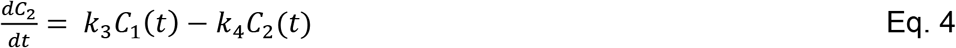

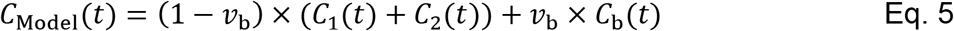

*C*_p_(*t*) is the input function, *C*_1_(*t*) and *C*_2_(*t*) are the concentrations in the first and second tissue compartments. *K*_1_ [mL/cm^3^/min] and *k*_2_ [1/min] represent the uptake and clearance rates from the plasma to the first tissue compartment, whereas *k*_3_ [1/min] and *k*_4_ [1/min] are the rate constants describing the exchange between the first and second tissue compartment (**Figure 3A**). *C*_Model_(*t*) shows the operational model curve, with the concentration *C*_b_(*t*) in the whole blood, and the blood volume fraction *v*_b_, which was set to 0.13, as previously reported for the myocardium [26]. Tissue distribution volume (*V*_T_) was calculated from the fit *K*_1_ and apparent rate constants *k*_2_, as well as *k*_3_ and *k*_4_ as follows: *V*_T_ = *K*_1_/*k*_2_(1+*k*_3_/*k*_4_).*V*_T_ was additionally assessed by graphical Logan analysis [27].

### Statistics

Continuous variables are presented as mean ± standard error of the mean (SEM). Student’s t-test and analysis of variance (ANOVA) tests were used for group comparisons of continuous variables. For multiple comparisons, Bonferroni correction was applied. Strength and direction of associations were assessed by Spearman’s rank-order correlation (r_s_). Outliers were assessed based on the Grubb’s test [28]. A two-tailed p-value of ≤ 0.05 was deemed statistically significant. Statistical analyses were carried out with SPSS (SPSS Statistics for Windows Version 24.0. IBM Corp. Armonk, NY).

## Results

### Rest/stress myocardial perfusion imaging

Time-activity curves (TACs) of ^18^F-flurpiridaz in the myocardium are depicted in **Figure 1**. At initial ^18^F-flurpiridaz injection, a rapid myocardial tracer inflow was observed, followed by a slow washout. Preliminary experiments revealed that a scan time of 40 min was sufficient to obtain accurate fits for the terminal washout phase (**Figure 1A**, model curve), thus allowing to account for residual myocardial activity from the initial ^18^F-flurpiridaz injection. Details on model curve fitting are given in the supplementary information. Subtraction of the extrapolated model curves from respective native TACs at 40-80 min post injection and correction for the injected ^18^F-flurpiridaz dose yielded the final stress TACs (**Figure 1B**). Notably, myocardial TACs were higher for the stress than for the rest scans at every measured time point, indicating a sustained increase of myocardial ^18^F-flurpiridaz uptake following regadenoson injection (**Figure 1B**). The increase of myocardial ^18^F-flurpiridaz uptake under pharmacological stress conditions was visually confirmed by PET images of the mouse myocardium (**Figure 2**).

**Figure 2:**
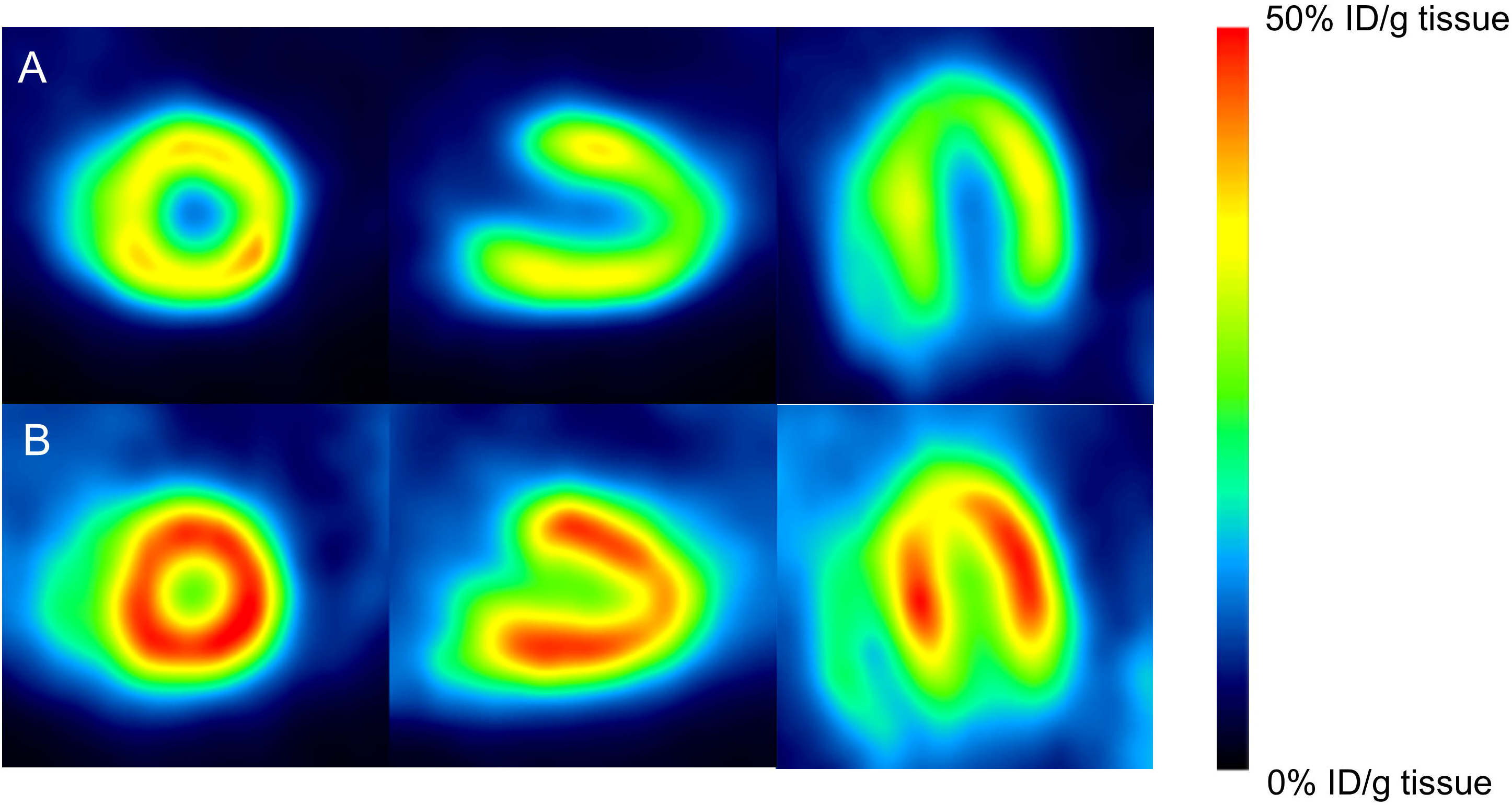
Positron emission tomography (PET) images of the mouse myocardium after ^18^F-flurpiridaz administration. **A**. Representative myocardial perfusion images at resting state. **B**. Representative myocardial perfusion images following regadenoson administration.

### Quantification of absolute myocardial blood flow by kinetic modeling

To determine absolute MBF, we employed kinetic tracer modeling using a metabolite-corrected input function that was recorded by continuous arterial blood sampling. Results of kinetic modeling were compared for one- and two-tissue compartment models. While the kinetic results were satisfactory by visual inspection for the two-tissue compartment model (**Figure 3**), the one tissue-compartment model did not provide accurate fitting (data not shown). *K*_*1*_, which is typically considered a measure of absolute MBF for tracers with high myocardial extraction fraction such as ^18^F-flurpiridaz [9], ranged from 6.7-20.0 mL/cm^3^/min, while myocardial volumes of distribution (*V*_*T*_) between 34.6 and 83.6 mL/cm^3^ were obtained. A representative example of the two-tissue compartment model fit is shown in **Figure 3B**. V_T_ values obtained from Logan graphical analysis were in agreement with the two-tissue compartment model fit (**Figure 3C**). These results suggest that the two-tissue compartment model constitutes a suitable model for the assessment of ^18^F-flurpiridaz kinetic parameters in mice, which is in agreement with previous studies in pigs and humans [21, 29, 30].

**Figure 3:**
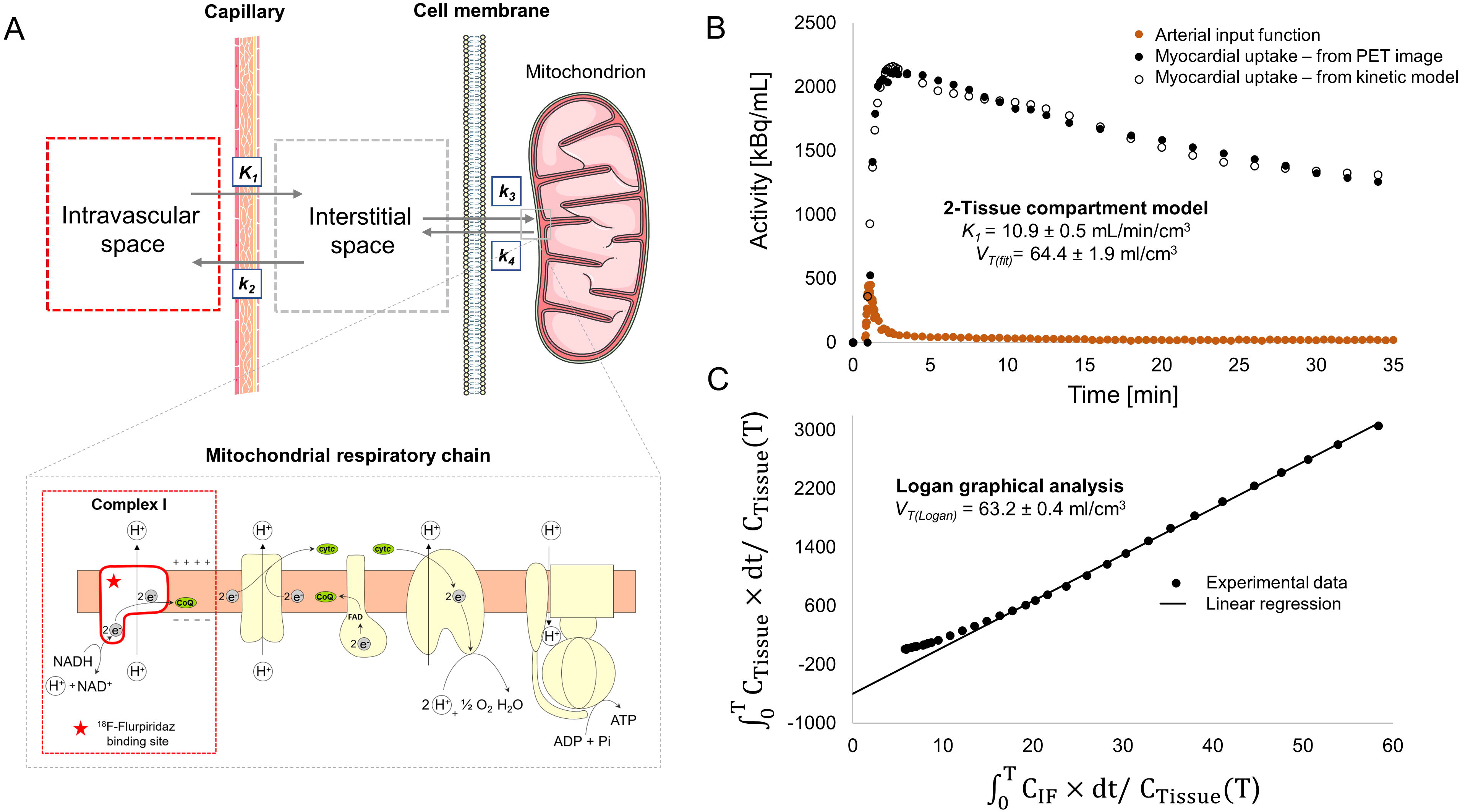
Two-tissue compartment model for ^18^F-flurpiridaz in the mouse myocardium. **A**. Visualization of the model. While *K*_*1*_ and *k*_*2*_ describe tracer delivery to the myocardium and clearance from the myocardium, respectively, *k*_*3*_ and *k*_*4*_ represent the association and dissociation rate constants towards mitochondrial complex I (MC-I), which is located at the mitochondrial membrane of cardiomyocytes. **B**. Representative two-tissue compartment fitting for the assessment of *K*_*1*_ and tissue volume of distribution (*V*_*T*_). **C**. Representative logan graphical analysis.

### Correlation of myocardial ^18^F-flurpiridaz uptake with *K*_*1*_ and *V*_*T*_

In a next step, we sought to assess whether the image-derived myocardial tissue uptake of ^18^F-flurpiridaz (%ID/g tissue), can be used as a surrogate measure for relative myocardial perfusion changes. Myocardial ^18^F-flurpiridaz uptake was significantly correlated with *K*_*1*_ from two-tissue compartment modeling. As shown in **Tables 1** and **2**, Spearman’s coefficients (r_s_) ranged between 0.478 and 0.573 for the rest scans and were higher after regadenoson injection (0.617-0.672), whereas significant correlations were obtained for all tested time slots. Nonetheless, R^2^ values were generally low, ranging from 0.21-0.43, which indicates that *K*_*1*_ does not explain a large proportion of the variability in image-derived myocardial ^18^F-flurpiridaz uptake.

**Table 1:**
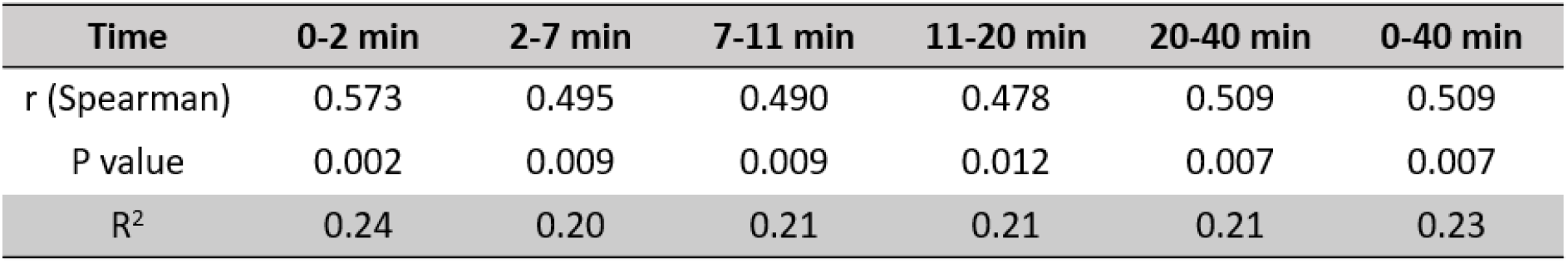
Correlation of image-derived myocardial ^18^F-flurpiridaz uptake with perfusion constant *K*_*1*_ at resting state (n=27).

| Time | 0-2 min | 2-7 min | 7-11 min | 11-20 min | 20-40 min | 0-40 min |
| --- | --- | --- | --- | --- | --- | --- |
| r (Spearman) | 0.573 | 0.495 | 0.490 | 0.478 | 0.509 | 0.509 |
| P value | 0.002 | 0.009 | 0.009 | 0.012 | 0.007 | 0.007 |
| R<sup>2</sup> | 0.24 | 0.20 | 0.21 | 0.21 | 0.21 | 0.23 |

**Table 2:** Correlation of image-derived myocardial ^18^F-flurpiridaz uptake with perfusion constant *K*_*1*_ following regadenoson (n=27).

| Time | 0-2 min | 2-7 min | 7-11 min | 11-20 min | 20-40 min | 0-40 min |
| --- | --- | --- | --- | --- | --- | --- |
| r (Spearman) | 0.672 | 0.681 | 0.676 | 0.671 | 0.617 | 0.668 |
| P value | <0.001 | <0.001 | <0.001 | <0.001 | <0.001 | <0.001 |
| R<sup>2</sup> | 0.43 | 0.42 | 0.41 | 0.38 | 0.33 | 0.41 |

In contrast, when myocardial ^18^F-flurpiridaz uptake was compared to the respective volumes of distribution (*V*_*T*_), excellent correlations were obtained between %ID/g tissue and *V*_*T*_ at rest (**Figure 4**). Notably, it was found that the two-tissue compartment model described the myocardial ^18^F-flurpiridaz uptake most accurately at later time frames (rest scans: **Figure 4C**, 7-11 min post injection, R^2^=0.98, **Figure 4D**, 11-20 min post injection, R^2^=0.99 and **Figure 4E**, 20-40 min post injection, R^2^=0.99), while earlier time frames were associated with lower R^2^ values (**Figure 4A**, 0-2 min post injection; **Figure 4B**, R^2^=0.71, 2-7 min post injection, R^2^=0.95. Similarly, the correlation model between %ID/g tissue and *V*_*T*_ was superior at later time frames following regadenoson injection, as depicted in **Figure 5**. Overall, %ID/g tissue, averaged from 20-40 min post injection, was most accurately described by the two-tissue compartment model under rest conditions and following pharmacological stress testing.

**Figure 4:**
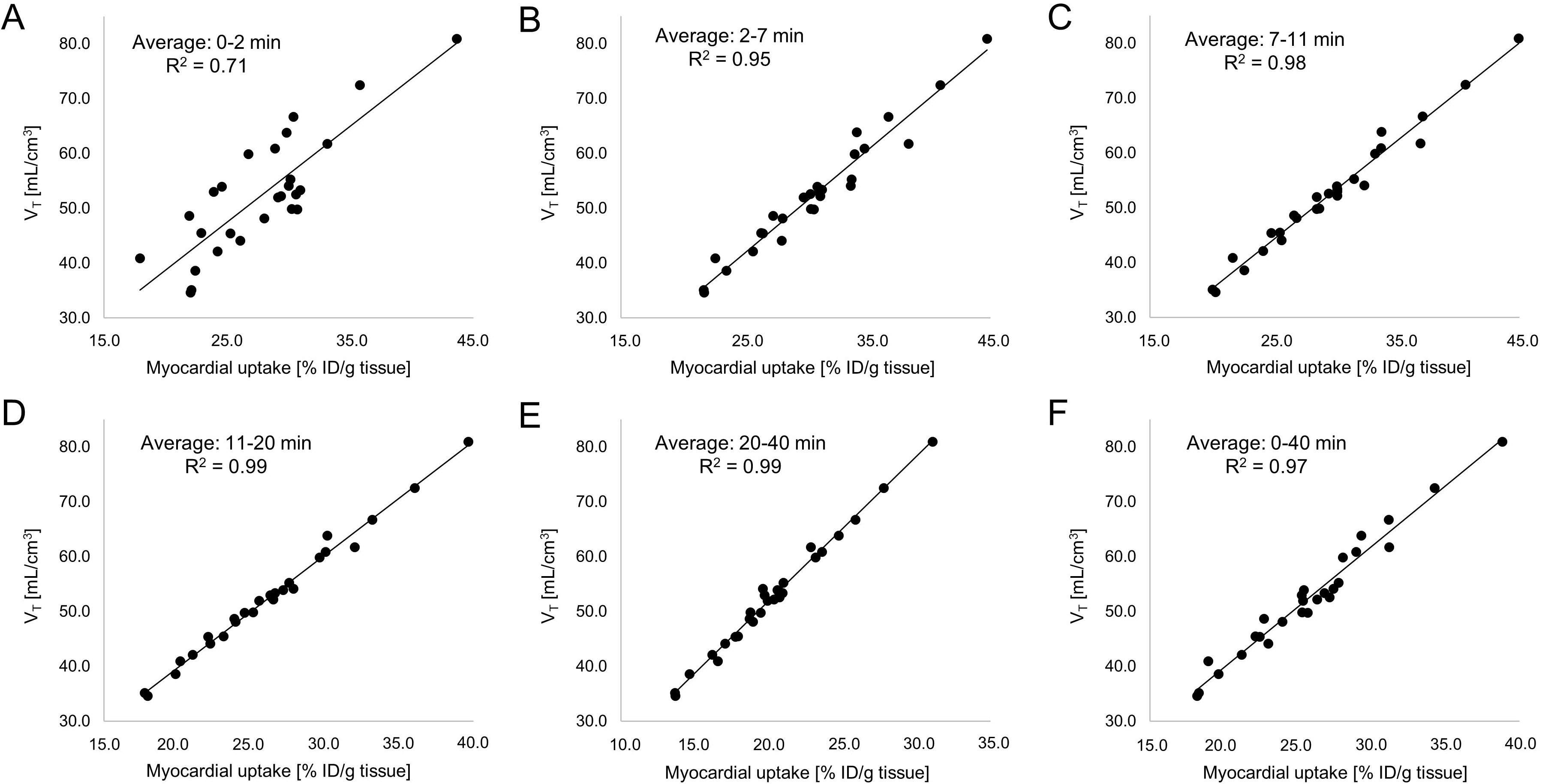
Resting state (no regadenoson) correlations of image-derived myocardial ^18^F-flurpiridaz uptake (%ID/g tissue), averaged at different time intervals, with tissue volume of distribution (*V*_*T*_). Early time intervals revealed lower R2 values, while later time intervals reached an R^2^ value of up to 0.99, indicating an excellent correlation between *V*_*T*_ and %ID/g tissue ^18^F-flurpiridaz. Averaged time intervals included (**A**) 0-2 min, (**B**) 2-7 min, (**C**), 7-11 min, (**D**) 11-20 min, (**E**) 20-40 min and (**F**) 0-40 min post injection.

**Figure 5:**
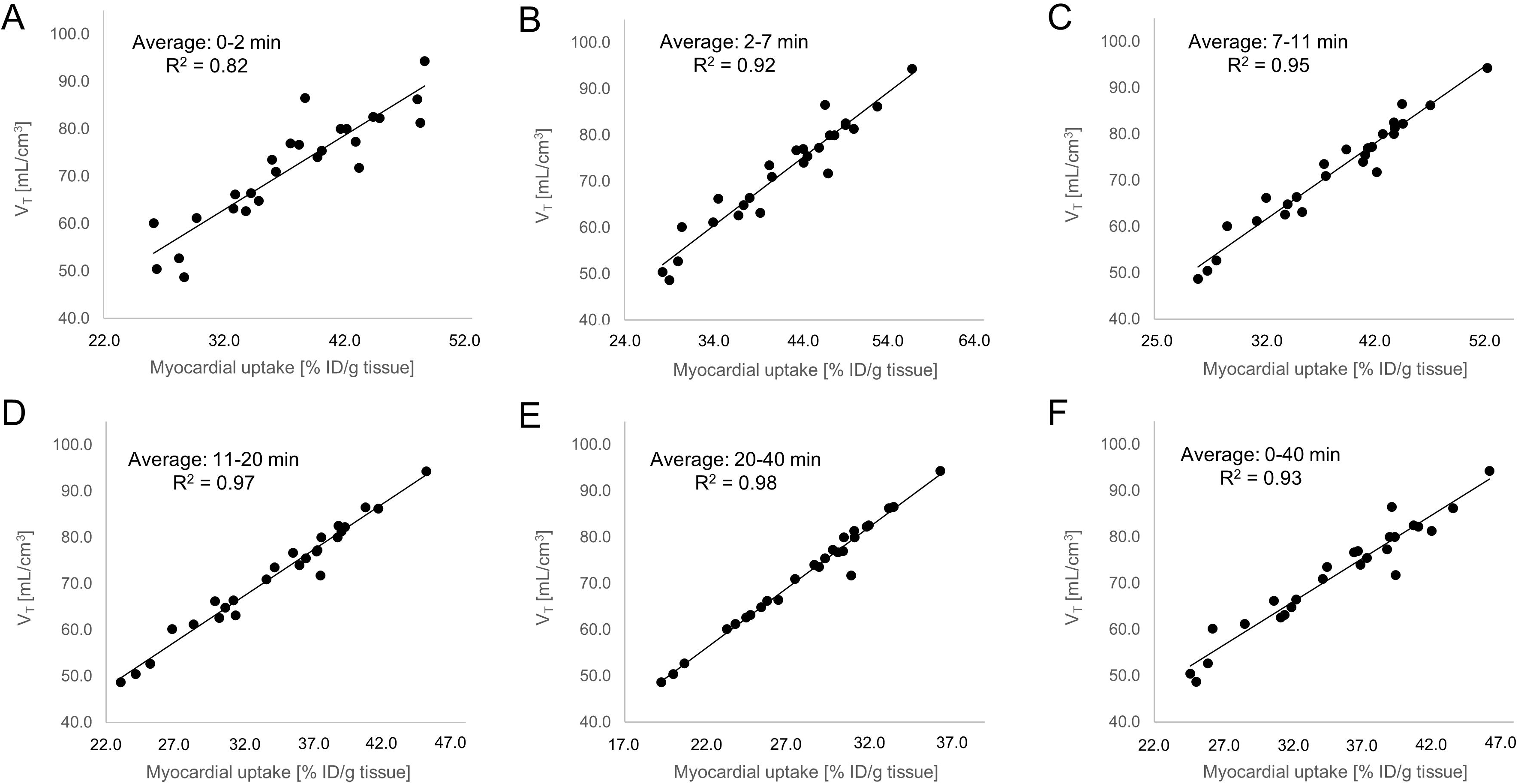
Correlations of image-derived myocardial ^18^F-flurpiridaz uptake (%ID/g tissue) following regadenoson-induced stress, with tissue volume of distribution (*V*_*T*_). Early time intervals revealed lower R^2^ values, while later time intervals reached an R^2^ value of up to 0.98, indicating an excellent correlation between *V*_*T*_ and %ID/g tissue ^18^F-flurpiridaz. Averaged time intervals included (**A**) 0-2 min, (**B**) 2-7 min, (**C**), 7-11 min, (**D**) 11-20 min, (**E**) 20-40 min and (**F**) 0-40 min post injection.

### Coronary Flow Reserve

CFR can be calculated from the ratio of MBF during regadenoson induced vasodilation to the MBF under resting conditions [31]. Given that *K*_*1*_ constitutes a measure of absolute MBF, we used the ratio of *K*_*1*_ following regadenoson injection to *K*_*1*_ at resting condition to calculate the CFR. As expected, *K*_*1*_ was significantly higher following regadenoson injection (**Figure 6A**, p<0.001). Similarly, regadenoson-induced coronary vasodilation led to a significant increase in *V*_*T*_ (**Figure 6B**, p<0.001). The resulting mean stress-to-rest ratios for *K*_*1*_ and *V*_*T*_ were identical (**Figure 6C**, 1.41±0.29 vs 1.41±0.17, p=0.92), however, with a higher variability obtained for *K*_*1*_. These results support the concept that both, *K*_*1*_ and *V*_*T*_, are similarly sensitive to pharmacologically induced changes of MBF in mice, while *V*_*T*_ seems to constitute a more robust parameter for the estimation of CFR. Notably, it was found that stress-to-rest ratios of image-derived myocardial ^18^F-flurpiridaz uptake accurately predicted CFR, as depicted in **Figure 7**. While tracer washout led to a gradual reduction of averaged %ID/g tissue over time, under resting conditions (**Figure 7A**) and following regadenoson injection (**Figure 7B**), the stress-to-rest ratio did not differ significantly between time intervals (**Figure 8C**). With a mean stress-to-rest ratio of 1.40±0.20, %ID/g ^18^F-flurpiridaz from 20-40 min post injection provided the most accurate mean ratio, as compared to CFR determined from *K*_*1*_ by the two-tissue compartment model.

**Figure 6:**
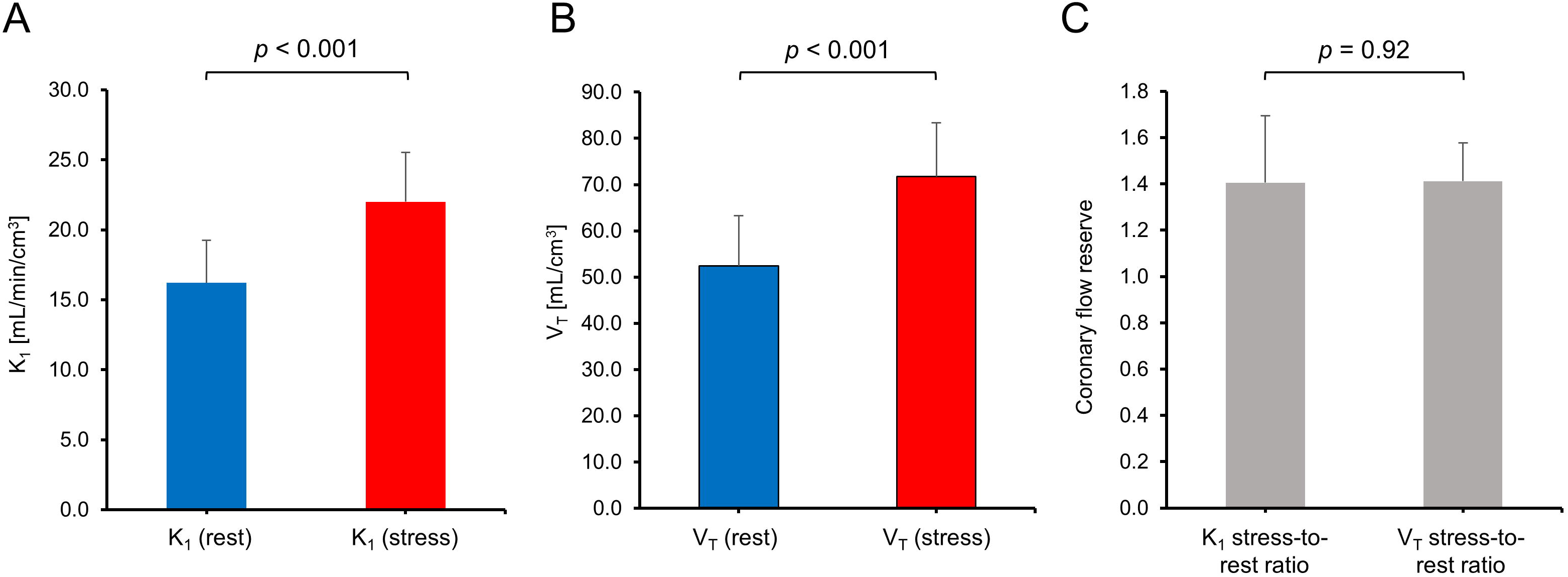
Average *K*_*1*_ and *V*_*T*_ under resting conditions and following pharmacological stress testing. **A**. *K*_*1*_ was significantly higher following regadenoson administration (p<0.001). **B**. *V*_*T*_ was significantly elevated following regadenoson administration (p<0.001). **C**. Mean coronary flow reserve, estimated from stress-to-rest ratios of *K*_*1*_ and *V*_*T*_.

**Figure 7:**
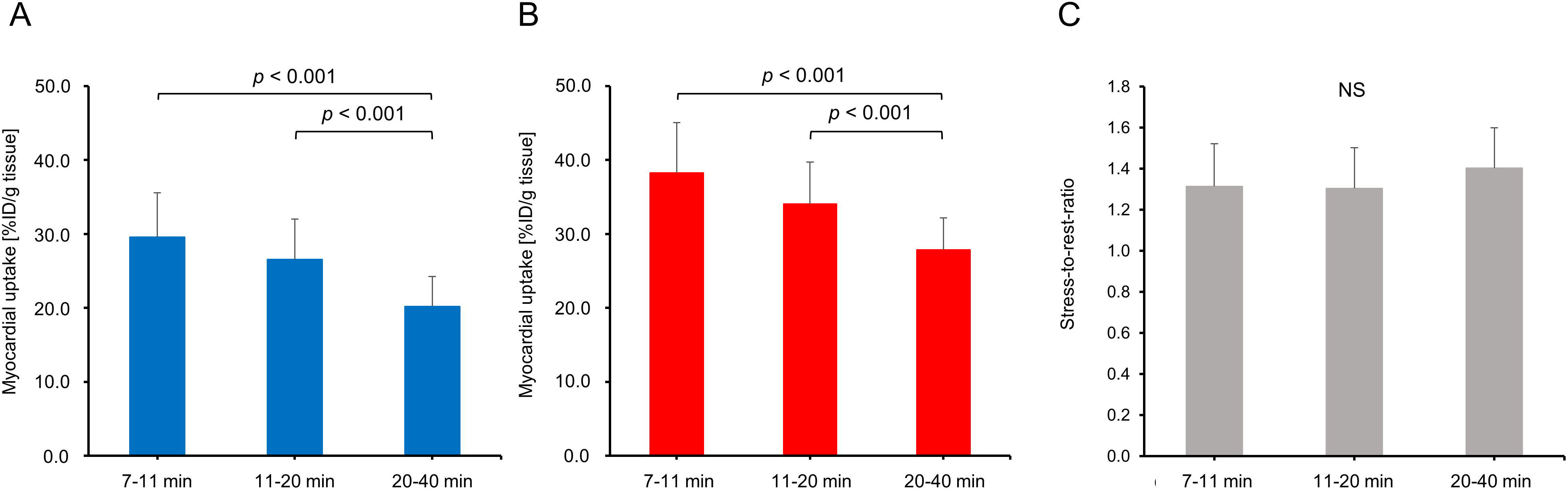
Myocardial ^18^F-flurpiridaz uptake averaged from 7-11, 11-20 and 20-40 min post injection. A significant reduction of averaged %ID/g tissue of ^18^F-flurpiridaz was found towards later time intervals (**A**) at resting state as well as (**B**) following pharmacological stress testing. **C**. In contrast, stress-to-rest ratios did not significantly differ between different time intervals.

## Discussion

In the present study, we show that rest/stress MPI with ^18^F-flurpiridaz in mice follows a two-tissue compartment model. While *K*_*1*_ and *V*_*T*_ were both similarly sensitive to vasodilator-induced increase of MBF, *V*_*T*_ exhibited a superior correlation with image-derived myocardial tracer uptake at all measured time intervals. In particular, image-derived tracer uptake at later time intervals revealed ideal coefficients of determination (R^2^ up to 0.99) when correlated with *V*_*T*_. Further, the average stress-to-rest ratio of myocardial tracer uptake (%ID/g tissue), obtained from 20-40 min post injection, was identical with the stress-to-rest ratios of *K*_*1*_ and *V*_*T*_. These results suggest that myocardial ^18^F-flurpiridaz uptake (%ID/g) in mice, as determined by from dynamic PET, correlates with MBF and can be used to estimate relative myocardial perfusion changes as well as CFR when pharmacological stress testing is employed.

To the best of our knowledge, this is the first study to assess the feasibility of rest/stress PET-MPI with ^18^F-flurpiridaz in mice. In agreement with previous reports in other species, including pigs [6] and humans [30], we found that a two-tissue compartment model was suitable to predict ^18^F-flurpiridaz uptake in the mouse myocardium. Nonetheless, significant species-differences were observed with regard to tracer washout from the myocardium. While myocardial ^18^F-flurpiridaz washout was negligible in pigs and humans, our time-activity curves unveiled a demonstrably faster kinetics in mice with evident washout from the mouse myocardium starting at 5-10 min post injection [6, 30]. The latter points towards species-differences in myocardial tracer retention that need to be taken into consideration for kinetic modeling. As such, while *k*_*4*_ (dissociation rate constant) was manually set to zero in previous studies with other species [6, 21, 32], we did not employ any restrictions for *k*_*4*_ in mice. In contrast to the two-tissue compartment model, the one-tissue compartment model did not result in satisfactory results by visual inspection, suggesting that binding to MC-I has a significant contribution to the kinetics as well as perfusion-linked ^18^F-flurpiridaz delivery to the mouse myocardium.

Despite the favorable properties of ^18^F-flurpiridaz for PET-MPI, the sustained myocardial radioactivity from the initial scan may affect the time-activity curves upon the second injection – particularly if rest and stress scans are performed on the same day. In our study, pilot experiments were conducted to determine the minimum scan duration required to allow adequate extrapolation of the terminal phase via non-linear regression models. Employing these models, we were able to correct for the residual myocardial radioactivity from the first ^18^F-flurpiridaz injection. In light of the observed species differences in ^18^F-flurpiridaz clearance from the myocardium, we conclude that a careful evaluation of the terminal phase for the underlying animal species seems crucial for 1-day rest/stress PET-MPI protocols with ^18^F-flurpiridaz.

Absolute MBF quantification has proven its utility in clinical routine, however, the latter requires an arterial input function [33] that can be derived from the PET image by placing a VOI in the left ventricular cavity [34]. While image-derived input functions are routinely obtained in clinical PET-MPI, this approach is hampered in mice due to the relatively small left ventricular dimensions. Accordingly, quantification of absolute MBF in mice requires arterial blood sampling, which is associated with significant technical challenges. To circumvent this limitation, we assessed whether myocardial ^18^F-flurpiridaz uptake can be used to provide information on myocardial perfusion changes as well as fundamental kinetic parameters such as *K*_*1*_ and *V*_*T*_. Indeed, although correlations of *K*_*1*_ with image-derived myocardial ^18^F-flurpiridaz uptake were significant, R^2^ values were generally below 0.5, indicating a poor fit for all time frames investigated. In contrast, ideal fits were obtained for the correlation of *V*_*T*_ with myocardial ^18^F-flurpiridaz uptake, particularly at later time intervals such as 20-40 min post injection. Although *K*_*1*_ is closely linked to perfusion, our results suggest that the rapid tracer kinetics renders myocardial ^18^F-flurpiridaz uptake more highly weighted towards MC-I binding than absolute MBF in mice. Accordingly, the relation between *K*_*1*_ and myocardial ^18^F-flurpiridaz uptake did not allow the prediction of absolute values for MBF from myocardial ^18^F-flurpiridaz uptake in our study. Notably, however, %ID/g ^18^F-flurpiridaz was highly correlated with *V*_*T*_. Indeed, our findings suggested that *V*_*T*_ was reliably estimated from myocardial ^18^F-flurpiridaz uptake in the absence of an arterial input function. Given (1) that *V*_*T*_ is sensitive to changes in *K*_*1*_ according to the equation *V*_*T*_ = *K*_*1*_ * (1+k_3_/k_4_)/k_2_ and (2) that mean rest-to-stress ratios were comparable for *V*_*T*_ and *K*_*1*_, it seems that changes in myocardial perfusion were accurately reflected by the model microparameter *V*_*T*_. Accordingly, although %ID/g ^18^F-flurpiridaz was not suitable to estimate values for absolute MBF, our data indicates that %ID/g ^18^F-flurpiridaz can be used for intra- and interindividual comparisons of relative myocardial perfusion in mice. Further, mean CFR values were comparable when calculated from stress-to-rest ratios of *K*_*1*_, *V*_*T*_ and %ID/g ^18^F-flurpiridaz from 20-40 min post injection, suggesting that stress-to-rest ratios of %ID/g ^18^F-flurpiridaz can be used as a surrogate of CFR when pharmacological stress testing is employed. In contrast to pharmacological stress testing, however, it should be noted that the assessment of CFR from exercise stress testing is not recommended in the absence of an input function, particularly due to the substantial increase of cardiac output under exercise testing, which leads to a proportional dilution of the intravenous radiotracer dose [35, 36].

There are study limitations that should be pointed out. First, high isoflurane concentrations can prompt coronary vasodilation and may have affected outcome measures of our study [37]. However, previous work has shown that heart rate and mean systolic blood pressure remain stable at isoflurane concentration up to 2.0% in mice [38, 39]. To minimize the vasodilatory properties of isoflurane, its concentration was therefore kept at or below 2.0% throughout the entire experiment. Second, an experimental arterial input function was only available in five animals and the average of all five input functions was used as a population (surrogate) input function for kinetic modeling in animals without arterial shunt, as previously reported [40].

In conclusion, our results indicate that ^18^F-flurpiridaz can be used for rest/stress myocardial perfusion imaging in mice and follows a two-tissue compartment model. Myocardial ^18^F-flurpiridaz uptake was equally sensitive to vasodilator-stress as *K*_*1*_ and *V*_*T*_, suggesting that a simplified assessment of CFR, based on image-derived findings, is feasible when pharmacological stress testing is employed. Further, although arterial blood sampling is required for absolute quantification of MBF, myocardial ^18^F-flurpiridaz uptake can be used for relative comparison of myocardial perfusion in mice.

## Compliance with ethical standards

### Funding

CG was supported by grants from the Swiss National Science Foundation (SNSF), the Olga Mayenfisch Foundation, Switzerland, the OPO Foundation, Switzerland, the Novartis Foundation, Switzerland, the Swiss Heart Foundation, the Helmut Horten Foundation, Switzerland, the EMDO Foundation, Switzerland, and the Iten-Kohaut Foundation, Switzerland. SB was supported by the University of Zurich (UZH) Foundation and by the Swissheart Foundation. AH was supported by the Swiss National Science Foundation (SNSF) and the University of Zurich (UZH) Foundation.

